# k-Nearest Neighbour Adaptive Sampling (kNN-AS), a Simple Tool to Efficiently Explore Conformational Space

**DOI:** 10.1101/2025.03.05.641673

**Authors:** Evianne Rovers, Anvith Thudi, Chris Maddison, Matthieu Schapira

## Abstract

Molecular dynamics (MD) simulations are computationally expensive, a limiting factor when simulating biomolecular systems. Adaptive sampling approaches can accelerate the exploration of conformational space by running repeated short MD simulations from well-chosen starting points. Existing approaches to adaptive sampling have been optimized to either guide sampling in a desired direction or explore well-formed convex spaces. Here, we describe a novel adaptive sampling algorithm that leverages a k-nearest neighbour (k-NN) graph of the sampled conformational space to preferentially launch explorations from boundary states. We term this approach k-Nearest Neighbor Adaptive Sampling (kNN-AS) and show it has state-of-the-art performance on simple and complex artificial energy functions and generalizes well on a protein test case. Implementation of kNN-AS is light, simple and better suited to complex real-world applications where the dimension and shape of the energy landscape is unknown.

## Introduction

Molecular dynamics (MD) simulations are used to understand the dynamics of biomolecular systems, which in turn can help to investigate the molecular mechanism of disease-related mutations^1–6^, uncover cryptic ligand binding sites^7–11^, or predict how allosteric compounds induce conformational shifts^12,13^. MD simulations are computationally intensive, and large systems often fail to reach equilibrium, let alone explore the full space of conformational states. Indeed, simulating processes such as protein folding, or protein complex formation, occur on timescales that can require months or even years of GPU time.

Methods have emerged that seek to explore the free energy landscape of a system more efficiently and can be roughly divided into adaptive and enhanced sampling approaches^14^. Enhanced sampling approaches, like simulated annealing and metadynamics, alter the energy of the entire system, to try and accelerate exploration of conformational space^14^. However, for large molecular systems these methods become computationally resource intensive^14^. Also, heating a multiprotein system can result in disassembly of the complex rather than conformational searching as seen for PROTAC-induced ternary complexes^15^. Adaptive sampling, in contrast, seeks to accelerate exploration of conformational space under normal conditions^16^. Here, long MD-simulations are divided into repeated short simulations, where starting states are carefully selected to optimize for exploration. Algorithms that select initial states seek to balance exploration and exploitation of the free-energy landscape defined by a feature set, termed collective variables (CVs). The results of these simulations are then typically combined and analyzed for long time-scale dynamics with Markov State Models^17^. Overall, adaptive sampling results are obtained under normal conditions, and thus do not require distribution reweighing^16^, however choice of algorithm is key to ensure efficient conformational search.

Where a specific goal in the conformational space is sought, the gradient of a feature / CV can be used to drive exploration. The algorithms FAST^17^ and AdaptiveBandit^18^ both excel at this task. However, these methods cannot be used when the desired direction or space of the exploration are unknown. In these latter cases, REAP/MAREAP use reinforcement learning to learn efficient exploration directions in the feature/CVs space^19^. It treats features/CVs independently and rewards states that result in CVs that deviate furthest from the population of known states^19^. The MAREAP algorithm was shown to outperform Adaptivebandit in a protein conformational sampling task^19^. Nonetheless, this method performs best with linear landscapes where there is only one optimal direction of exploration or when the direction switches from one CV to another (for example, a L-shaped landscape). Multi-agent TSLC, a different algorithm, performs better with non-linear landscapes^19^. Since the shape of the lansdcape is typically unknown, which algorithm is most appropriate *a piori* is unclear. Therefore, an ideal algorithm would efficiently explore the conformational space regardless of dimension and shape.

In this application note, we present a simple algorithm that leverages a k-Nearest Neighbor (k-NN) graph applied to a set of known molecular states and identifies edges of the conformational space for focused sampling. We term this approach k-Nearest Neighbor Adaptive Sampling (kNN-AS). This approach builds on extensive work showing the power of k-NN graphs for searching complex feature spaces, for example, as employed by Google’s PageRank algorithm^20^. We briefly describe the algorithm and then demonstrate that it achieves superior performance on both simple and complex artificial energy functions, as well as a real-world protein MD simulation case. The code for the algorithm and experiments can be found on github: https://github.com/ERovers/kNN-AS

## Algorithm

One goal of adaptive sampling is to select starting states which maximize the number of new states observed during the simulation. The key challenge is finding a local heuristic that globally optimizes this goal. Reinforcement learning (RL) is one approach, but this can introduce modeling biases and/or require many simulations to learn an adequate model. We instead develop a simple heuristic based on our observations of molecular dynamics simulations: we found that the average distance and direction of a state to its nearest neighbours (amongst seen states) is an effective indicator of how many new states the dynamics will explore.

Inspired by this, our algorithm uses a k-NN approach to construct a graph from a point cloud (states in feature/CV space). The edges that connect a vertex (state) to its k-NNs are summed to create a directional vector. New points for the next generation of simulations are then selected based on the magnitude of this vector (Algorithm 1). Intuitively, this algorithm promotes exploration of states at the boundary of a free energy landscape (Figure 1).

**Algorithm 1.**
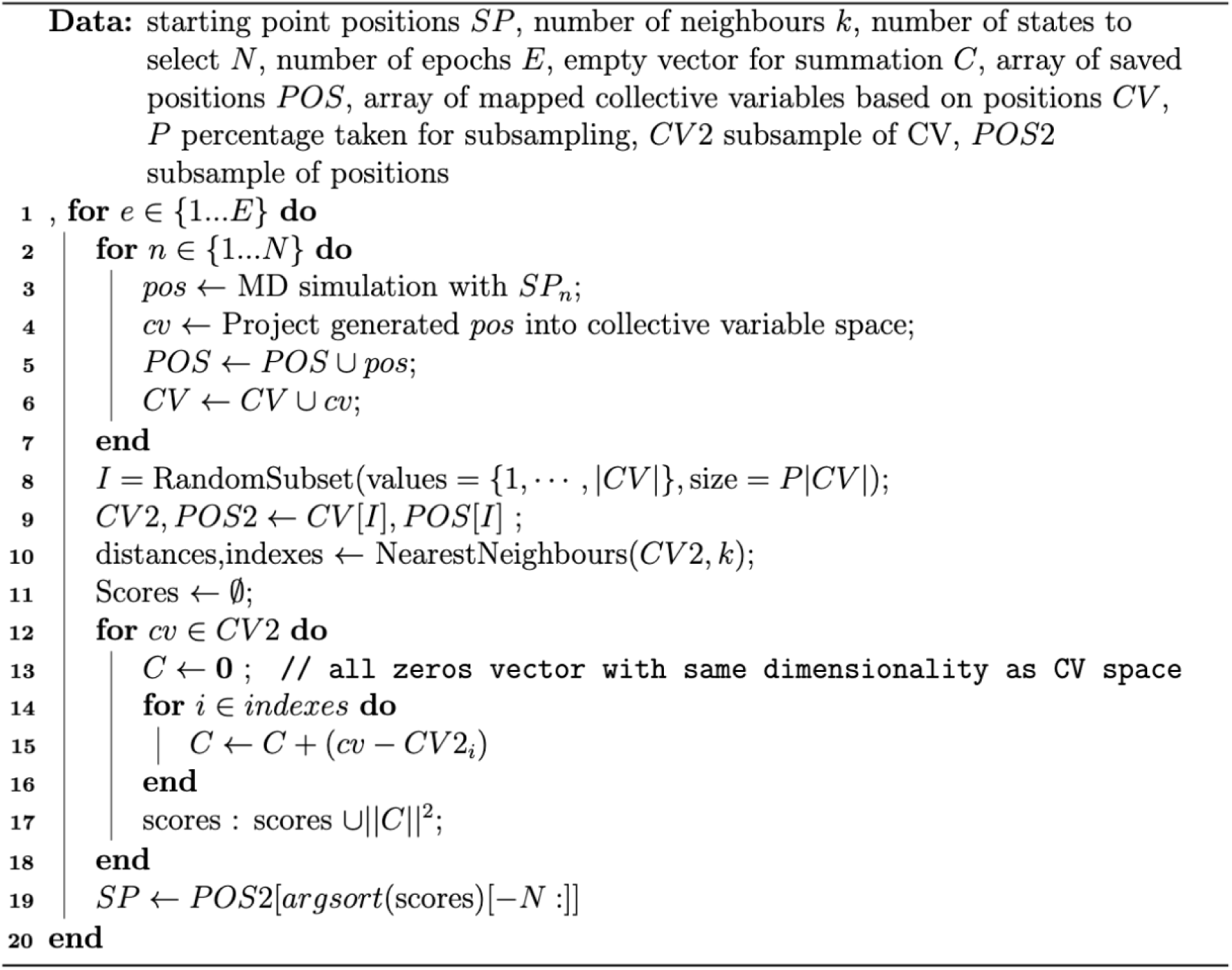
kNN-AS sampling algorithm to determine states at the boundary of a free energy landscape.

**Figure 1.**
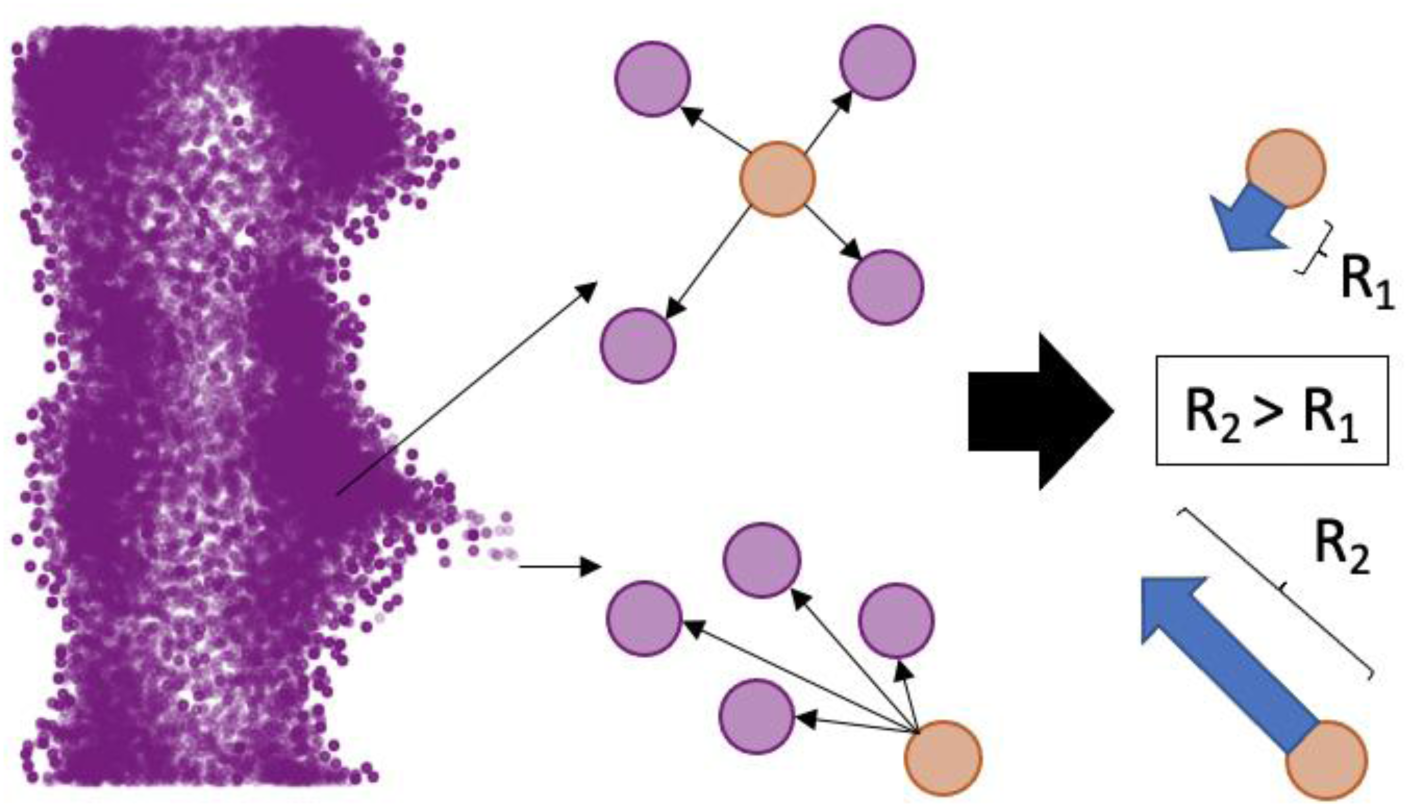
Visualization of the algorithm. Datapoints within in point cloud have neighbours on all sides which results in an overall small directional vector (R_1_) and datapoints on the edge will have neighbours concentrated on one side resulting in large directional vector (R_2_). Edges can be selected based on the size of the overall directional vectors.

The kNN-AS algorithm has two hyperparameters specific to the algorithm: the size of subsample (P) used for generating the k-NN graph, and number of neighbours (k). When performing a test study on an artificial energy landscape (Figure 2A), we found that the size of subsample is the most important hyperparameter, since the extent of exploration increases when the size of the subsample decreases (Figure 2B). The reason for this behaviour could be that subsampling results in a neighbourhood structure of the k-NN graph that reflects the global structure of a point cloud, as opposed to the local structure induced by the autocorrelations in MD trajectories when using the whole dataset. In particular, during MD simulations, frames are saved every few timesteps. If this interval is too short, the frames can form a nearly continuous line in the conformational landscape, leading k-NN graph to pick neighbours from the same trajectory and causing the graph to only detect local structure. To check for autocorrelation, a global set of angles between the vertices can be calculated which should show a uniform distribution (Figure 3, top right). If a skewed distribution is observed with peaks around 0 and 180 degrees (Figure 3, top left), then k-NN graph only detects the local structure of the MD trajectory itself, namely all points form a line (Figure 3, bottom left). Subsampling addresses this by removing the neighbouring frames, creating a sparser and more random dataset. This results in selecting states on the edges of the explored space (Figure 3, bottom right), which further promotes exploration.

**Figure 2.**
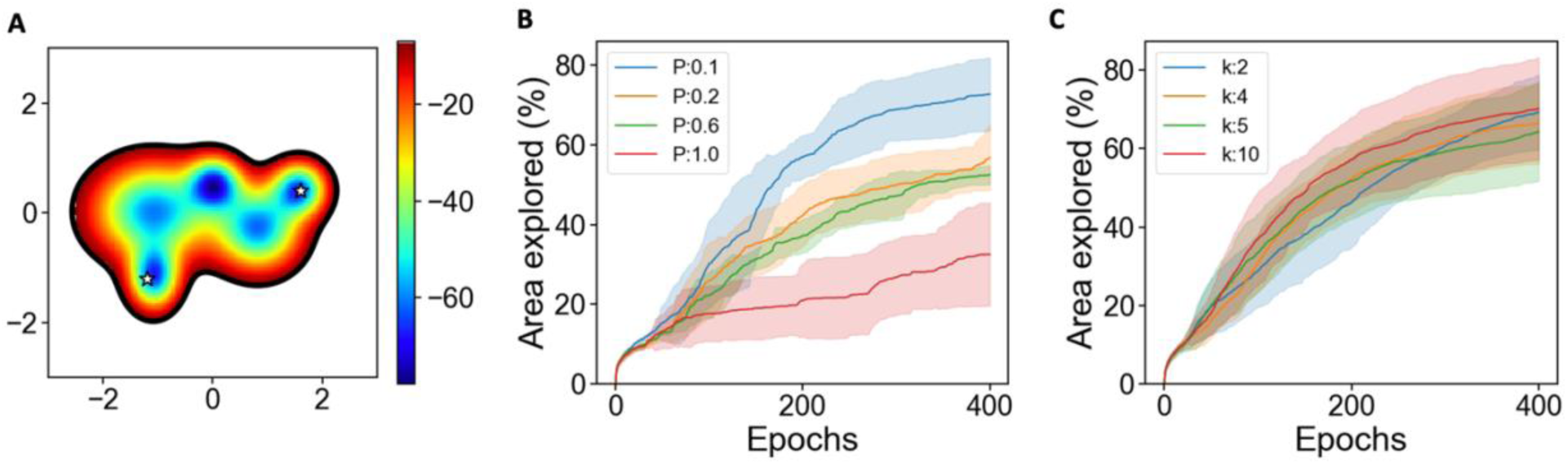
The effect of hyperparameters on exploration of artificial landscape. A) Artificial energy landscape with multiple local minima and one global minimum. B) Decreasing the size of subsample while keeping a fixed number of neighbours (k=5) leads to more extended exploration (9 replicate). C) Altering the number of neighbours when fixing subsample size (P=0.1) has no effect on the number of visited states (9 replicates).

**Figure 3.**
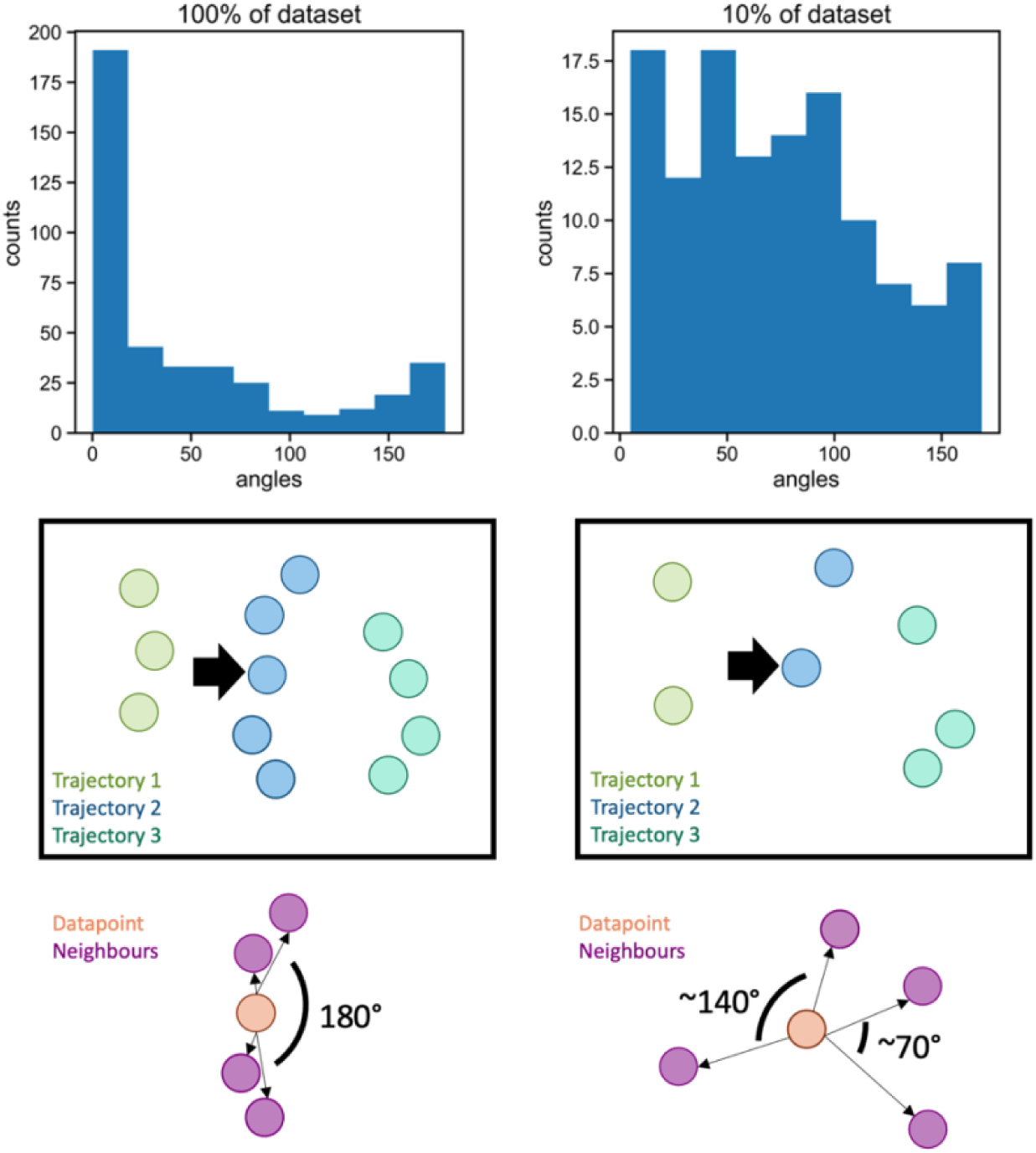
Effect of subsampling on kNN graph and state selection. Using the whole dataset (left), the four neighbours of the datapoint (indicated with the black arrow) are from the same trajectory and the overall distribution between the neighbour vertices are skewed to 0 and 180. When selecting a subset of the data, the four neighbours for the selected datapoint are from different trajectories. This results in a uniform distribution of angles between neighbour vertices.

The other hyperparameter, number of neighbours (k), has no clear effect on the total exploration (Figure 2C). When visualizing the states selected, we do see that a high value of k favors states at the outer boundary of the explored space, whereas a low value of k also selects states bordering “local” edges or open patches within the explored space, for example the gap between the small and big point cloud (Figure 4). This can be compared with the UMAP algorithm where a high number of neighbours extracts the global structure of the data while a low number concentrates on the local structure and finer details of the data^21^. Nonetheless, the number of neighbors has a subtle impact on total exploration, suggesting that variations in the number of neighbors likely only affect local exploration.

**Figure 4.**
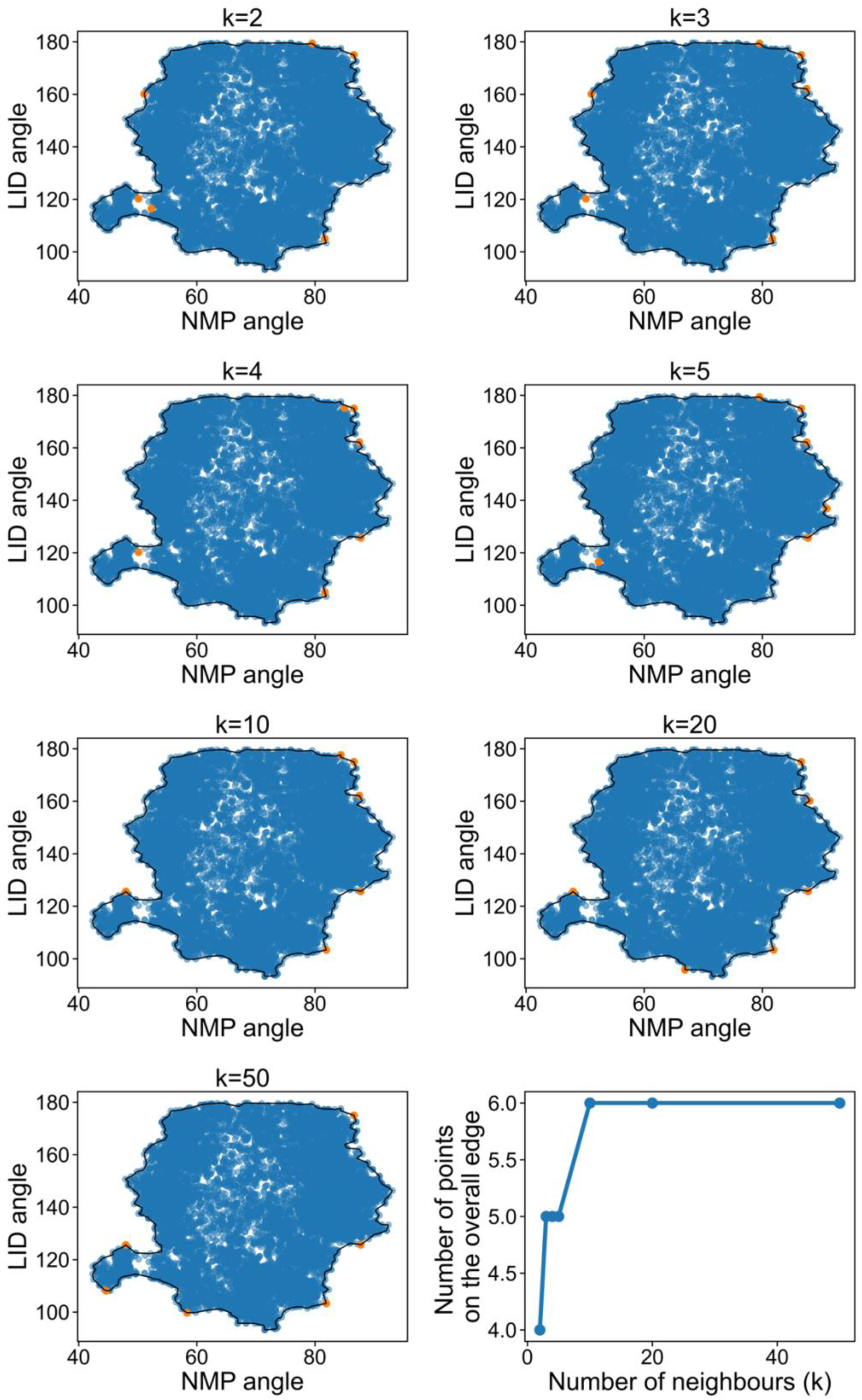
Differences in MD starting state selection when choosing 2, 3, 4, 5, 10, 20 or 50 neighbours. Explored states are shown in blue. Starting states for the next round of MD simulations are shown in orange. Last panel: Number of selected starting states not on the outer edge of the explored space.

Other hyperparameters, such as number of MD simulations per epoch, number of timesteps per simulation, and the number of epochs, are universal to any adaptive sampling strategy, and are highly dependent on the computer resources available and the system that is explored. For example, the power of adaptive sampling approach is that MD simulations can be run in parallel, and therefore, higher number of simulations per generation is better as more area can be explored within the same wall-clock time. However, the realistic number of simulations is set by the number of resources available. Additionally, each individual trajectory must be long enough to capture new states while remaining short enough to take advantage of an adaptive sampling strategy. Lastly, the number of epochs can be determined based on a convergence criterion, for example, the extent of the area explored. For more consideration on these hyperparameters, we would refer to reviews and other adaptive sampling papers and review^16,18,19,22–24^.

## Results

### Artificial energy potentials

To test how well kNN-AS performs compared with MAREAP, considered the state-of-the-art in using RL in adaptive sampling^19^, we first considered four simple artificial energy function test cases (Figure 5). In these test cases we used potentials where a single particle diffuses following a Langevin dynamic. The first two potentials are the symmetric cross potential where the energy is equal across all arms, and asymmetric cross potential where there is an energy well in one of the arms. These are key test-cases used to demonstrate the performance of MAREAP^19^. The simulations started on the horizontal arms of the energy landscape and performance was measured by calculating the area explored by each method. The explored area is calculated as the total number of grid cells (grid size is 0.05) divided by the number of cells visited at least once. Each simulation was performed 100 times to generate error bounds. For the kNN-AS algorithm the k was set to 5 and P to 0.1.

**Figure 5.**
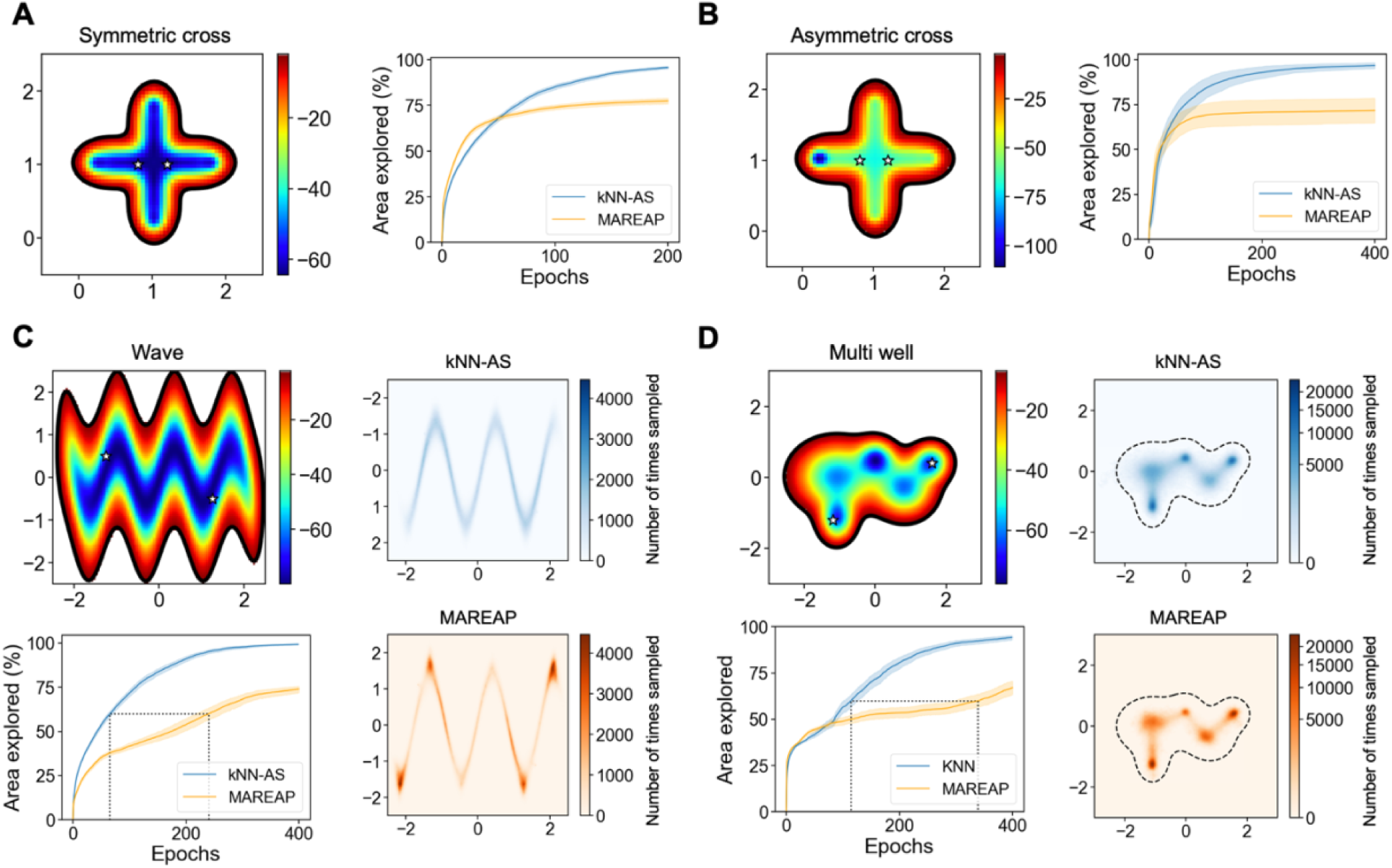
kNN-AS efficiently explores complex artificial potential functions. A) symmetric cross potential, B) asymmetric cross potential, left plots show visualization of the energy landscape with stars indicating starting locations and right plot shows the area explored for kNN-AS and MAREAP over time. C) wave potential, and D) multi well potential, the top left shows a visualization of the landscape with stars indicating the starting locations. Bottom left panels show the area explored over time with kNN-AS and MAREAP. The right-hand panels show the distribution of the landscape explored by both kNN-AS and MAREAP. The color bar indicates the number of times the state was sampled.

In the symmetric cross landscape, kNN-AS and MAREAP explored at similar speed for the first 50 epochs (Figure 5A), but MAREAP tends to jam at the extremity of the cross, leading to only 7 out of 100 replicates exploring 90% or more of the landscape. kNN-AS on the other hand explored 90% of the landscape in 89 out of 100 replicates. Similarly with the asymmetric cross landscape, MAREAP reached 90% exploration in 1 replicate where all replicates in kNN-AS explored 90% of the landscape or more (Figure 5B).

We also investigated the versatility of the kNN-AS algorithm with more complex non-linear landscapes. The wave potential is a landscape based on a *sin* wave function where the energy is equal across the wave. The landscape is considered complex due to the quick directional changes. MAREAP explored most of the landscape without getting trapped in corners, hence the exploration curve did not plateau (Figure 5C). However, plotting state distributions reveals that simulations over-sampled the top and bottom of the wave near the starting points (Figure 5C). kNN-AS exploration was more efficient, with 60% of the landscape visited within 66 epochs versus 240 epochs on average for MAREAP. Since the energy in this system is uniform across the wave, it does not reflect real-life test cases where energy minima can be separated by high-energy transition states. To test a more realistic energy landscape, we generated a multi-well landscape with multiple local minima and one global minimum with higher energy transitions connecting the lower energy states. Here, MAREAP often found the local minima adjacent to the starting points but failed to find the global minimum in 64 out of 100 replicates. Also, only seventeen replicates visited more than 90% of the landscape (Figure 5D). kNN-AS always found the global minimum, but still did not always explore the whole landscape as only 69 replicates visited more than 90% of the landscape. Overall, these artificial potential functions showed that kNN-AS explores linear and complex energy landscapes efficiently and fast.

### Adenylate Kinase

Adenylate kinase is a protein consisting of three domains and is mostly studied for its transition pathway from closed to open state. This transition can be described by the angles between the smaller domains (LID (residues 30-78) and NMP (residues 116-164)) and the larger core domain. We set up an experiment by using the open and closed crystal structures as starting points (PDB: 4AKE^25^ and 1AKE^26^ respectively) and the angles as the feature/CVs for exploration. MAREAP and kNN-AS were run for 400 epochs in triplicate as described in the Methods section with k set to 5 and P to 0.5 for the kNN-AS algorithm. A continuous MD simulation was also run as a baseline. MAREAP and kNN-AS were both run for an aggregated simulation time of 240 ns (6 simulations of 0.1 ns per epoch for 400 epochs), half starting from the open kinase state and half from the closed kinase conformation.

After 240 ns, kNN-AS had explored on average 48% of the space, MAREAP 36% and full MD 20% (Figure 6A). This shows that both MAREAP and kNN-AS explored significantly more area than the baseline MD simulation. Interestingly, both MAREAP and kNN-AS retrieved an alternate crystal structure (PDB: 2AK3^27^) that was missed by the regular MD simulation (Figure 6B). To evaluate whether kNN-AS pushes the simulations to unrealistic conformations, we visualized the structures at the edges of the free energy landscape (Figure S1) and did not observe signs of irregularities such as alpha distorted helices/beta sheets or unfolding of protein regions.

**Figure 6.**
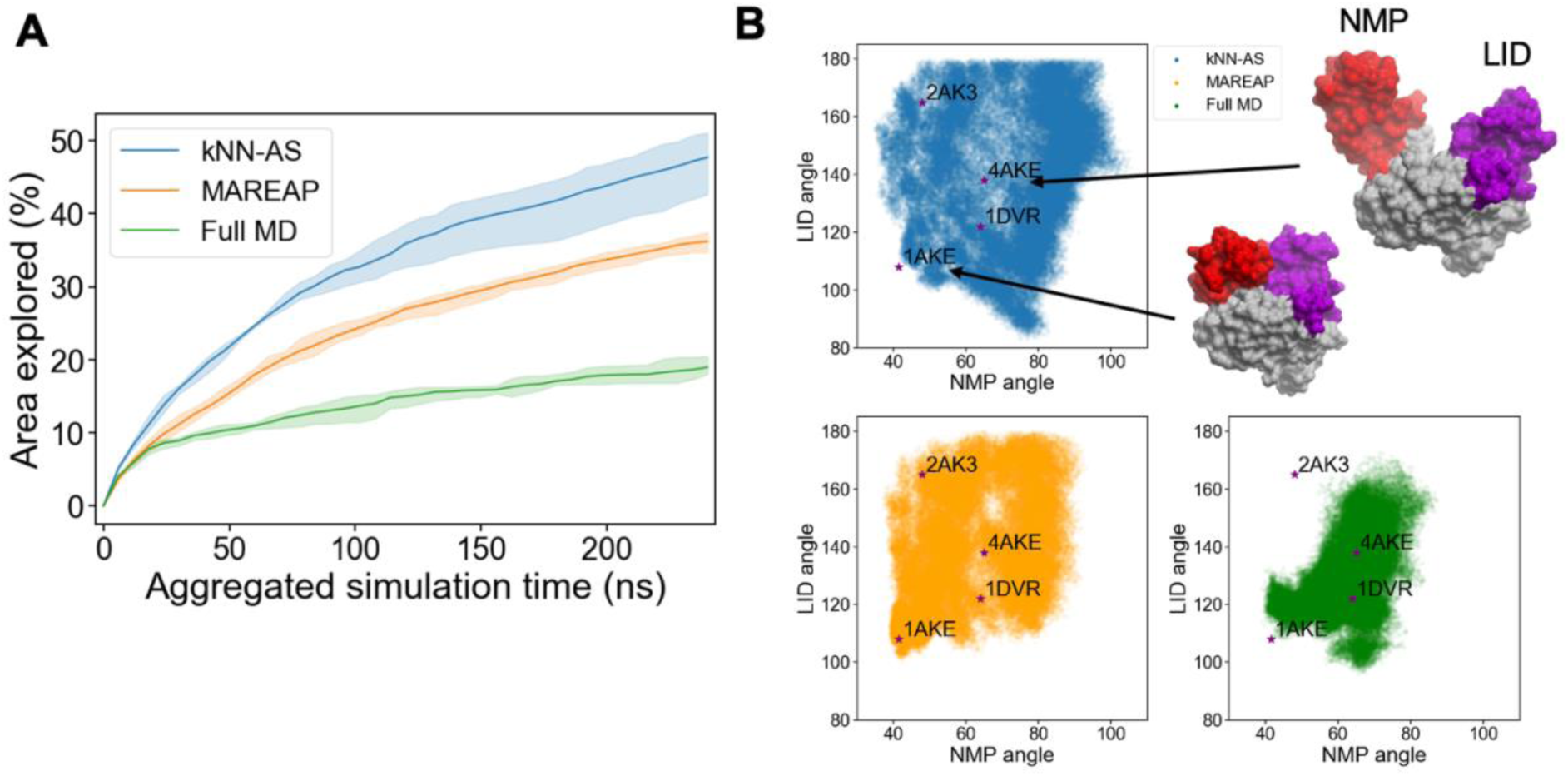
Results on Adenylate Kinase conformational landscape. A) Percentage of area explored vs epochs. B) Visualization of area explored in LID/NMP angle space for each method, when starting from open (PDB: 4AKE^25^).) and closed (PDB: 1AKE^26^) conformations. Experimentally observed alternate conformations are shown on the landscape (PDB: 2AK3^27^ and 1DVR^28^).

## Discussion

MD simulations of large biomolecular systems, such as multi-protein complexes, often require impractical amounts of resources to reach equilibrium. Therefore, optimized strategies to sample the free-energy landscape aim to accelerate this process by either running parallel simulations with carefully chosen starting states (adaptive sampling)^18,19,23,24^, or by introducing a bias to push the simulations towards underexplored areas or other regions of interest (enhanced sampling)^18,19,23,24^. The latter requires reweighing of the energy landscape^14^ while adaptive sampling methods are run under normal conditions and can be easily analyzed using Markov State Models for underlying dynamics^17^. Using artificial neural networks to learn quantum-mechanical force fields or rapidly generate ab initio protein equilibrium ensembles are also novel and promising avenues to accurately and efficiently sample conformational states^29–32^. In this application note, we present a very simple adaptive sampling strategy where a kNN-graph detects states on the edge of the conformational landscape and determines which areas have been explored.

Since kNN-AS is based on the kNN graph, it may suffer from the curse of dimensionality when using high-dimensional CV space^33^. Also, kNN-AS efficiently explores the space defined by a set of CVs, but it does not learn which CVs are most important for sampling, unlike MAREAP. Therefore, a future direction could be to pair kNN-AS with dimension reduction techniques, such as principal component analysis^34^, variational mixture of posteriors^35^ or variational autoencoders^36,37^ to extract a lower CVs space for more efficient exploration.

An important hyperparameter is the size of subsample used in constructing the kNN-graph. Using a subset introduces a level of randomness in the process allowing for heavily explored areas to be explored again if only a small set is sampled from the region. This can be advantageous for landscapes with high energy transition states, as MD simulations need to be restarted multiple times at the same edge before a transition is observed. Importantly, the size of subsample is dependent on the number of frames saved during the simulations. Shorter intervals cause frames to be too close in space, resulting in near neighbours always coming from the same trajectory and limiting efficient exploration across trajectories. Therefore, it is recommended to either use small data samples when generating k-NN graphs, or keeping long intervals between saved frames, or both. For the test cases shown here, subsamples of 10% and 50% of the data were used for the artificial energy potentials and the adenylate kinase cases respectively. The lower number of frames required for kNN-AS to perform well has benefits for simulating large biomolecular system as less data needs to be stored in memory.

The number of neighbours in kNN is a hyperparameter that influences state selection on local versus global edges (Figure 4), but has less of an effect on the total exploration. We decided to use five nearest neighbours, since this value of k showed good performance on the artificial landscapes, but future research could further interrogate how focus on local versus global exploration differently affects results of MD simulations. Since kNN-AS only requires hyperparameter tuning for 2 parameters, k and P (aside from regular adaptive sampling hyperparameters), a few combinations can be tested on shorter runs for a given system before launching longer simulations.

Overall, we have shown that kNN-AS is a simple and short algorithm that efficiently guides the exploration of complex conformational landscapes with minimal hyperparameter tuning. This simple code can easily be inserted within more detailed algorithms, can be augmented with complementary techniques such as scores re-weighting for directed searches, or combined with deep learning approaches such as VAEs.

## Methods

### Langevin Dynamics

Four artificial energy potential functions were chosen to compare the performance of kNN-AS with MAREAP. The first two potentials (symmetric cross and asymmetric cross) have been previously described and tested for the MAREAP algorithm^19^. The other two energy potentials were developed to mimic more complex energy landscapes (wave and multi well potentials). The energy potentials are defined as:

#### Symmetric cross potential^19^

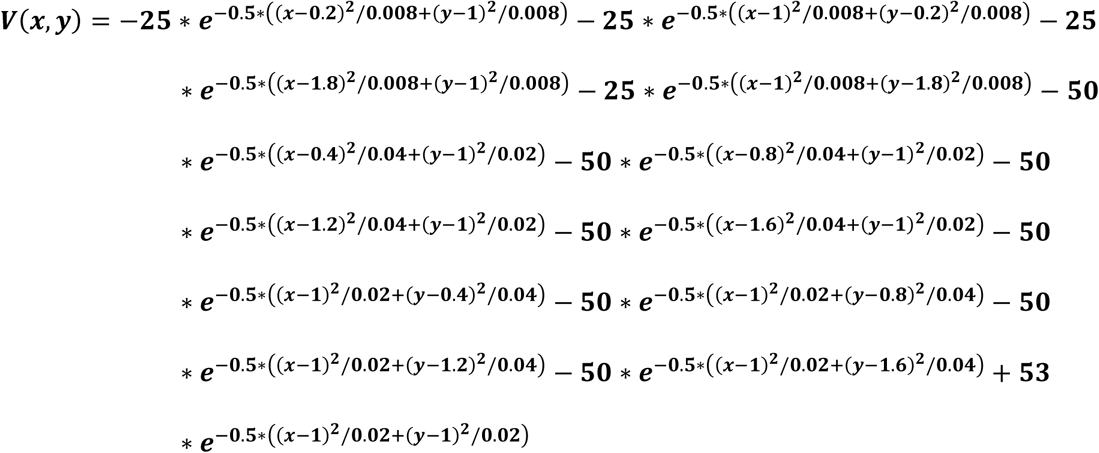

#### Asymmetric cross potential^19^

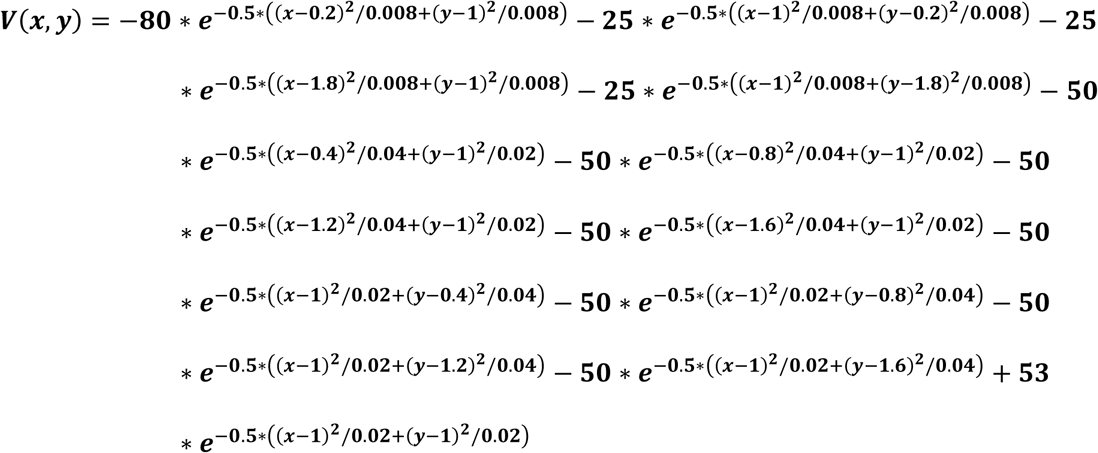

#### Wave potential

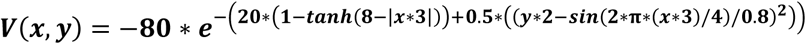

#### Multiwell potential

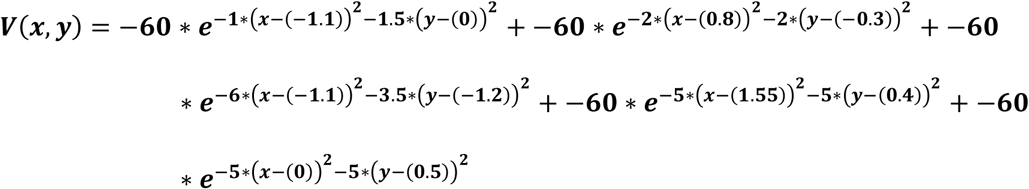

All simulations were run with OpenMM 7.7^38^ and a Langevin integrator. The timestep was set to 2 fs, the friction coefficient to 1 ps^-1^ and the temperature to 300K. The mass of the single particle was 100 Da. For both protocols, MAREAP and kNN-AS, 400 epochs were run with 6 MD simulations for each epoch (each running for 500 steps) and a frame was saved every 25 steps. Only the symmetric cross potential was run for 200 epochs. For each test case, 100 replicates were performed to determine error bounds. The starting locations are: symmetric cross [[0.8, 1], [1.2, 1]], asymmetric cross [[0.2, 1], [1.8, 1]], wave [[-1.25, 0.5], [1.25, -0.5]], and multi-well [[1.6, 0.4], [-1.2, -1.2]].

Hyperparameters for MAREAP were similar to the parameters previously described^19^ with 10^4^ frames used for κ-means clustering (with 5 restarts for random cluster initialization). The number of clusters was determined using the CLUST^39^ calculation by using d=2, γ=0.6 and b=3^-4^. The maximum weight change was set to δ=0.02, and collective rewards and fraction stakes were used. For kNN-AS, 10% of the data was subsampled for kNN graph generation and number of neighbours was set to 5.

### All-atom dynamics

MD simulations for Adenylate Kinase were also performed with OpenMM 7.7^38^ with a Langevin integrator, temperature of 300K, time step of 2 fs and friction coefficient of 1 ps^-1^. All bonds with hydrogens were constrained and the pressure was held at 1 bar using the Monte Carlo barostat. The starting states are the open and closed conformations (PDB: 4AKE^25^ and 1AKE^26^ respectively) and were prepared using the Amber 14 forcefield^40^ and TIP3-pfb^41^ water model. 40000 water molecules were added to the systems to obtain a similar number of atoms in both systems. The ionic strength was set to 0.1M. Then the system was energy-minimized and equilibrated for 100 steps.

MAREAP and kNN-AS were both run for 400 epochs with 6 MD simulations per epoch. Each MD simulation consisted of 50,000 steps and positions were saved every 3500 steps. To generate error-bounds, each method was run in triplicate. MAREAP was initialized using similar hyperparameters as the artificial energy function along with using 12 least-count clusters. kNN-AS was run with 5 neighbours for kNN graph generation and 50% of the data was used. Regular MD simulations were performed starting from the two starting points each for 120ns to get a total of 240ns aggregated simulation time, similar to MAREAP and kNN-AS.

## Acknowledgements

This research was enabled in part by support provided by Compute Ontario (https://www.computeontario.ca/) and the Digital Research Alliance of Canada (alliancecan.ca). M.S. and E.R. gratefully acknowledges support from NSERC [Grant RGPIN-2019-04416], CQDM (Quantum Leap-176), and MITACS accelerate (IT13051). The Structural Genomics Consortium is a registered charity (no: 1097737) that receives funds from Bayer AG, Boehringer Ingelheim, Bristol Myers Squibb, Genentech, Genome Canada through Ontario Genomics Institute [OGI-196], EU/EFPIA/OICR/McGill/KTH/Diamond Innovative Medicines Initiative 2 Joint Undertaking [EUbOPEN grant 875510], Janssen, Merck KGaA (aka EMD in Canada and US), Pfizer, and Takeda.

C.M. is supported in part by the Province of Ontario, the Government of Canada through CIFAR, and companies sponsoring the Vector Institute and the support of the Natural Sciences and Engineering Research Council of Canada (NSERC), RGPIN-2021-03445. A.T. is supported by a Vanier Fellowship from NSERC.

## Data availability

The scripts to reproduce the results presented and the script containing the kNN-AS algorithm is available at: https://github.com/ERovers/kNN-AS. Raw results are available on request.

