## Supplementary material for "k-Nearest Neighbour Adaptive Sampling (kNN-AS), a Simple Tool to Efficiently Explore Conformational Space": Figure S1

**Supplementary Figures**

Figure S1..........................................................................................................................2

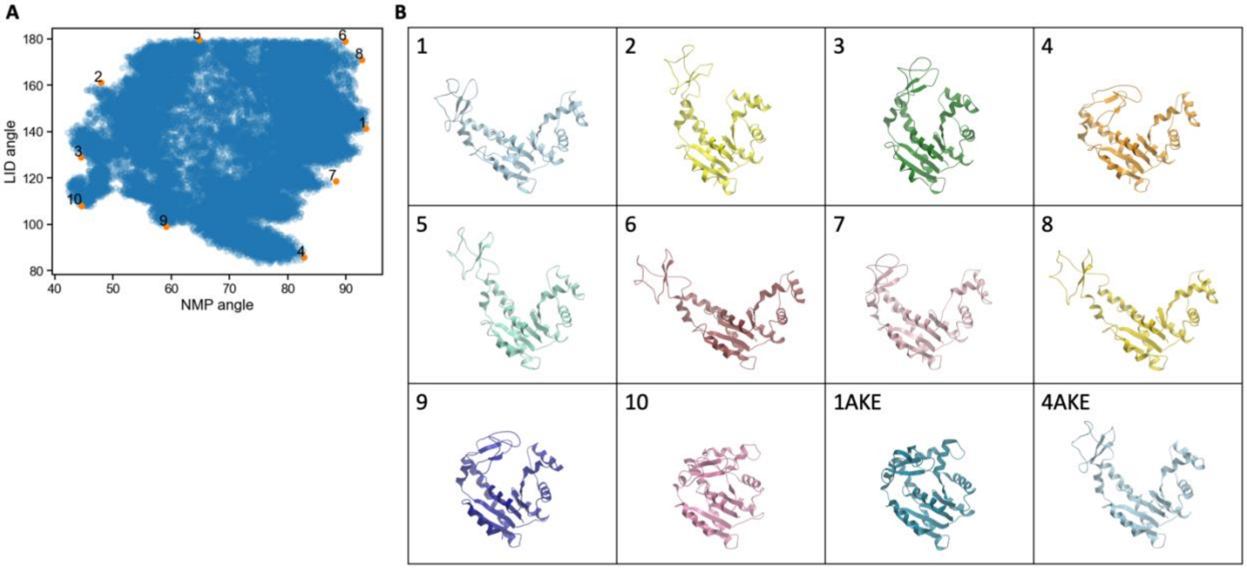


**Figure S1. Configurations of Adenylate Kinase at the edge of the conformational space.** A) Conformational landscape expressed in NMP and LID angle with states chosen at the edges shown and labelled in orange. B) 3D models of the selected states with corresponding labels. The crystal structures of the closed conformation (1AKE[26]) and open conformation (4AKE[25,26]) are shown as references.
